# T lymphocyte regulatory cytokines predict frailty in older adults

**DOI:** 10.64898/2026.04.16.716397

**Authors:** Thomas E Akie, Ethan Loew, Ziyuan Huang, Haley A Neff, Olivia P Michaels, John P Haran

## Abstract

Frailty is a multi-system syndrome causing increased susceptibility to health insults in older adults. Immune system dysregulation and inflammaging have emerged as mechanisms that may affect multiple organ systems in the frailty syndrome. This present study seeks to define the immune state in community-dwelling adults suffering from frailty. We evaluated a subgroup of 169 individuals enrolled in the Gut-brain Alzheimer’s disease Inflammation and Neurocognitive Study (GAINS). Participants in the GAINS study were scored for frailty using the Clinical Frail Scale. A panel of 27 inflammatory cytokines was analyzed from the serum of each participant. Frailty was present in 33 (19.5%) of the cohort, and was correlated with age, malnutrition, and cognitive assessments. Statistical analysis adjusting for clinical covariates revealed higher serum levels of IL-2, IL-10, and IL-17 in frail patients. Using machine learning classification, we developed a predictive model of frailty with strong discriminative performance (AUC 0.78). Individual element analysis via Shapley Additive Explanations (SHAP) revealed that inflammatory markers had the greatest influence on the model, and IL-7 was the single most important element in the prediction of frailty. Together, our data demonstrate a novel pattern in which T-cell regulatory inflammatory molecules as mediators of frailty, implicating cellular immunity as a potential mechanism of dysfunctional aging.

## INTRODUCTION

Aging is a progressive decline in molecular and physiological systems that is conserved across multiple levels of biology. Despite its inevitability, the process of aging is heterogeneous and not well understood^1^. Purported mechanisms of aging include mitochondrial dysfunction, genomic and epigenetic instability, loss of normal proteostasis, cellular senescence, and changes in intercellular communication^2^. Nevertheless, specific markers and targets of aging-related dysfunction in humans remain elusive.

Frailty is a syndrome of aging in older adults, which portends disproportionately adverse responses to physiologic stressors^3^. While a clinically distinct entity, frailty is thought to arise from dysregulation of processes implicated in aging at large – mitochondrial dysfunction and metabolic dysregulation, genetic and epigenetic alterations, and alterations in immune system function^4^. Hence, there is particular interest in understanding frailty as a means of understanding the underlying causes of disordered aging.

Disruptions to immune system function, or “inflammaging”, is of particular interest as a mediator of frailty since it is environmentally mediated and has the potential to communicate stress states between independent organ systems^5^. Subsets of frail patients have been found to have elevated levels of C-reactive protein^6^, IL-6^7^, and TNFα receptor^8,9^. Despite this, studies have failed to identify a specific pattern of inflammatory markers in frailty. Additionally, randomized trials investigating anti-inflammatory treatment have failed to prevent progression to frailty^10^. Thus, the precise role of peripheral inflammation in frailty has yet to be defined.

This study seeks to further define the role of whole-body inflammation in aging beyond traditional inflammatory markers. To do this, we investigated an expanded panel of immune regulatory factors and cytokines to determine the role of cellular immunity in frail older adults. Furthermore, we employed novel machine learning classification techniques to derive a predictive model of frailty using demographic and clinical factors and inflammatory markers.

## METHODS

### Study Setting and Patient Population

This study was a pre-planned subgroup analysis of the Gut-brain Alzheimer’s disease Inflammation and Neurocognition Study (GAINS). The complete details of this study are published elsewhere^11,12^. Briefly, community-dwelling adults aged 60 years or older were enrolled and underwent data and sample collection as follows. The GAINS study was approved by the institutional review board at the University of Massachusetts Chan Medical School, Worcester, MA.

### Data and Sample Collection

General demographic data, including age, sex, race, and medical history, were collected at the index visit. Patients underwent frailty, nutritional assessment, and cognitive screening at the index visit and at each subsequent visit. Frailty was primarily assessed using the Clinical Frail Scale (CFS)^13^, with a score of ≥4 deemed “frail”. The Edmonton Frail Scale has been used as a second means of frailty assessment to ensure accuracy of our primary frailty assessment^14^. Nutritional assessment was conducted using the Mini Nutritional Assessment (MNA)^15^.

Cognitive assessments were performed using the Alzheimer’s Disease Assessment Scale – Cognitive Subscale 13 (ADAS-Cog 13)^16^.

Venous phlebotomy was performed and peripheral blood collected in vacutainer tubes. Samples were centrifuged to obtain serum samples, aliquoted, and frozen at −80°C until analysis.

Demographic data, assessments, and tissue samples from the index visit were used for the present study.

### Inflammatory Marker Analysis

Absolute quantification of peripheral inflammatory markers was performed using the Bio-Plex Pro Human Cytokine 27-plex Assay (Bio-Rad), a multiplex screening panel enabling measurement of 27 cytokines, chemokines, and growth factors in a single sample. To prepare peripheral plasma samples for analysis, aliquots were thawed on ice, clarified by centrifugation to remove particulates, and diluted 1:4 in standard buffer. The manufactures recommended protocol was followed for bead incubation, detection antibody labeling, and signal amplification. Standards and quality controls supplied with the kit were included on each plate to generate calibration curves and assess assay performance across runs. Multiplex assay results were acquired on a Bio-Plex 200 system (Bio-Rad) equipped with Bio-Plex Manager software. Median fluorescence intensity (MFI) values for each analyte were converted to concentrations using 5-parameter logistic regression. Samples falling below the quantifiable range were assigned the lower or upper limit of detection. Final cytokine concentrations were used for downstream statistical analyses.

### Statistical Analysis

Demographics were analyzed using standard descriptive techniques. Where applicable, continuous variables are displayed using median ± interquartile range (IQR). Categorical variables are displayed as n (%).

For the purposes of analysis, frailty was treated as a binary categorical variable with CFS ≥ 4 being deemed frail^17^. Comparisons of demographics, clinical assessments, and individual inflammatory markers between groups were performed using Mann-Whitney tests, adjusting for multiple comparisons using the Holm-Šídák approach^18^. Correlation between CFS and the Edmonton Frail Scale was performed using linear regression.

For the adjusted analysis, covariates were assessed for multicollinearity using the variance inflation factor (VIF). Covariates with VIF > 5 were serially eliminated until no persistent collinearity was observed. Remaining covariates were selected for adjusted analysis using stepwise backward elimination (significance threshold of p < 0.2) to generate logistic regression model. Our final model included the following significant covariates: age, the presence of hypertension, use of antipsychotic medications, ADAS-Cog 13 Score, and MNA Score. Odds ratios for individual inflammatory markers were then adjusted using the logistic regression model. Adjustment for multiple comparisons was performed using the false discovery rate. We declared statistical significance at p<0.05 (after adjusting for multiple comparisons) and FDR < 0.05.

Machine learning classification was performed using a gradient-boosted decision tree algorithm (XGBoost)^19,20^ with Bayesian hyperparameter optimization embedded within a nested Monte Carlo cross-validation framework^21,22^. Five independent experiments, each with 10 splits, were performed using random seeds generated by seedhash based on the MD5 algorithm^23,24^. Experimental results were averaged with respect to final performance characteristics. To assess interactions among elements, Shapley Additive Explanations (SHAP) Analysis was performed^25–28^.

Analyses and figure generation were performed using Prism (GraphPad), R programming language (version 4.4.2, R Core Team, 2024), and Python (version 3.10.18, Python Software Foundation).

## RESULTS

### Frailty is associated with age, dementia, and malnutrition

We assessed the association of frailty with clinical comorbidities and markers of aging in a cohort of community-dwelling older adults. Clinical demographics of the cohort are displayed in Table 1. Frail patients were significantly older than their robust counterparts (Fig. 1a). As expected, patients deemed frail by the Clinical Frailty Scale also had significantly higher scores on the Edmonton Frailty Scale than robust patients (Fig. 1b), and these had a significant positive correlation (Supplemental Figure 1). Frailty was significantly associated with age, higher scores on the ADAS-Cog 13 assessment, and lower Mini-Nutritional Assessment scores (Fig 1c and 1d).

**Table 1.** Baseline characteristics of GAINS cohort.

|  | Robust<br>n = 136 | Frail<br>n = 33 |
| --- | --- | --- |
| Age, median [IQR] | 72 [66-76] | 76 [68-81] |
| Clinical Assessments, median [IQR] |  |  |
| Clinical Frailty Scale | 2 [1-3] | 4 [4-5] |
| Edmonton Frailty Scale | 1 [0-2] | 4.5 [2-6] |
| ADAS Cog-13 Score | 10.165 [6-14.415] | 22.33 [15-49] |
| Mini Nutritional Assessment | 26.5 [24-28] | 23 [19.5-26] |
| Charlson Comorbidity Index | 3 [2-5] | 4 [4-5] |
| Demographics, n (%) |  |  |
| Sex |  |  |
| Male | 52 (38.2) | 11 (23) |
| Female | 84 (61.8) | 22 (77) |
| Diabetes | 22 (16.1) | 5 (15.1) |
| Hypertension | 53 (38.9) | 21 (63.6) |
| Hyperlipidemia | 63 (46.3) | 13 (39.3) |
| Asthma | 16 (11.7) | 4 (12.1) |
| COPD | 3 (2.2) | 2 (6) |
| CKD | 4 (2.9) | 1 (3) |
| MI | 8 (5.8) | 3 (9) |
| CHF | 2 (1.4) | 0 (0) |
| Malignancy | 20 (14.7) | 5 (15.1) |
| Immunosuppression | 3 (2.2) | 2 (6) |
| Alzheimer's | 8 (5.8) | 10 (30.3) |
| Malnourished | 22 (16.6) | 17 (54.8) |
| Tobacco Use | 37 (27.2) | 10 (30.3) |

**Figure 1.**
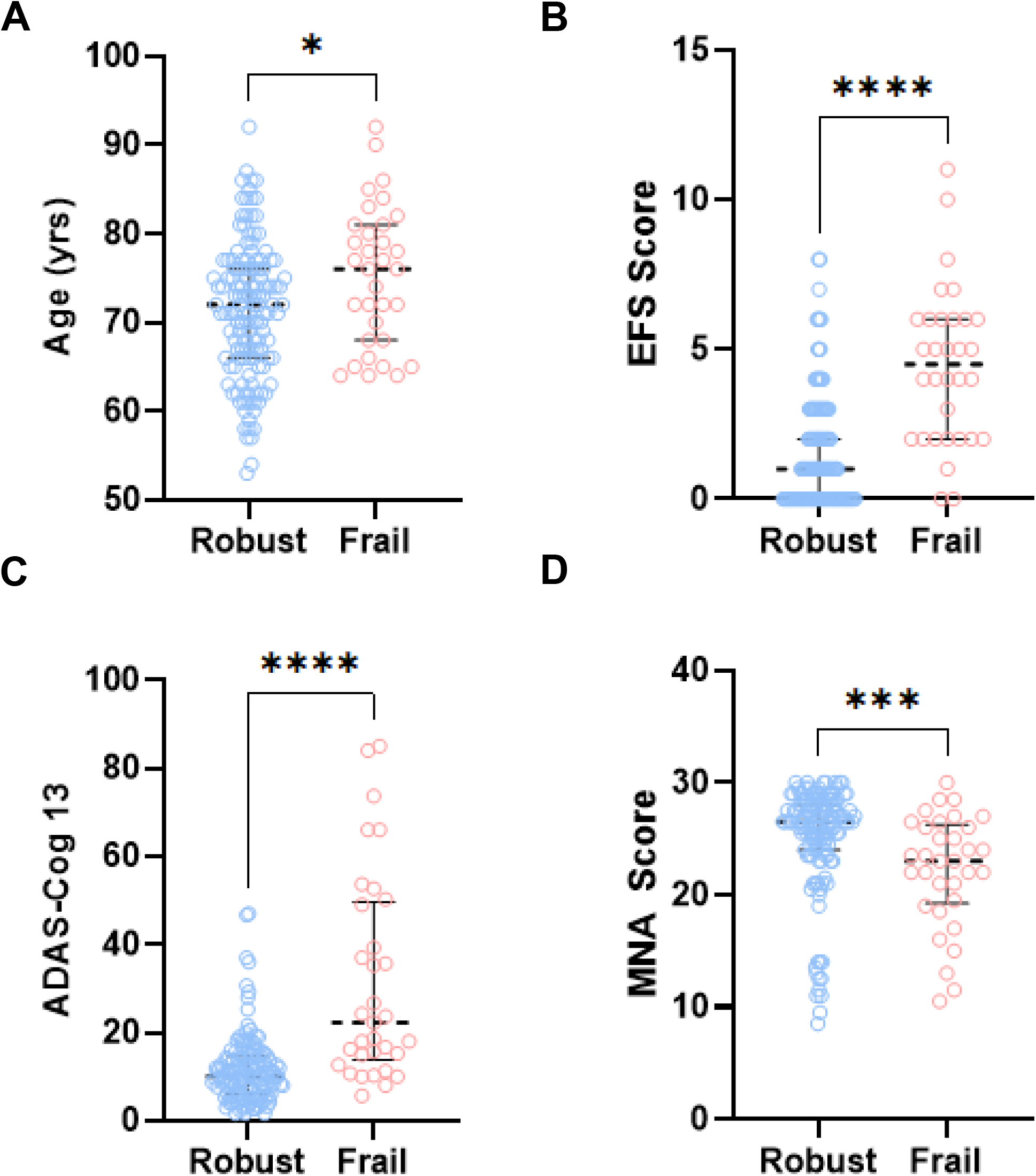
Comparison of Frail and Robust participants in GAINS cohort. Scatter plot of A) age, B) Edmonton Frailty Scale (EFS), C) Alzheimer’s Disease Assessment Scale–Cognitive Subscale (ADAS-Cog 13), and D) Mini Nutritional Assessment (MNA) between robust and frail participants in the GAINS cohort. Dotted line is median with interquartile range. *p < 0.05, ***p<0.005, ****p<0.0001.

### Pro-inflammatory cytokines are increased in frailty

To assess whether frailty is associated with increased whole-body inflammation, we first evaluated unadjusted levels of serum cytokines between robust and frail patients (Fig. 2). Frail patients demonstrated increases in several pro-inflammatory cytokines, with the most substantial significant differences demonstrated in IL-7 and TNFa. Additionally, IL-2, IL-10, and IL-17 were also found to be significantly different between the two populations.

**Figure 2.**
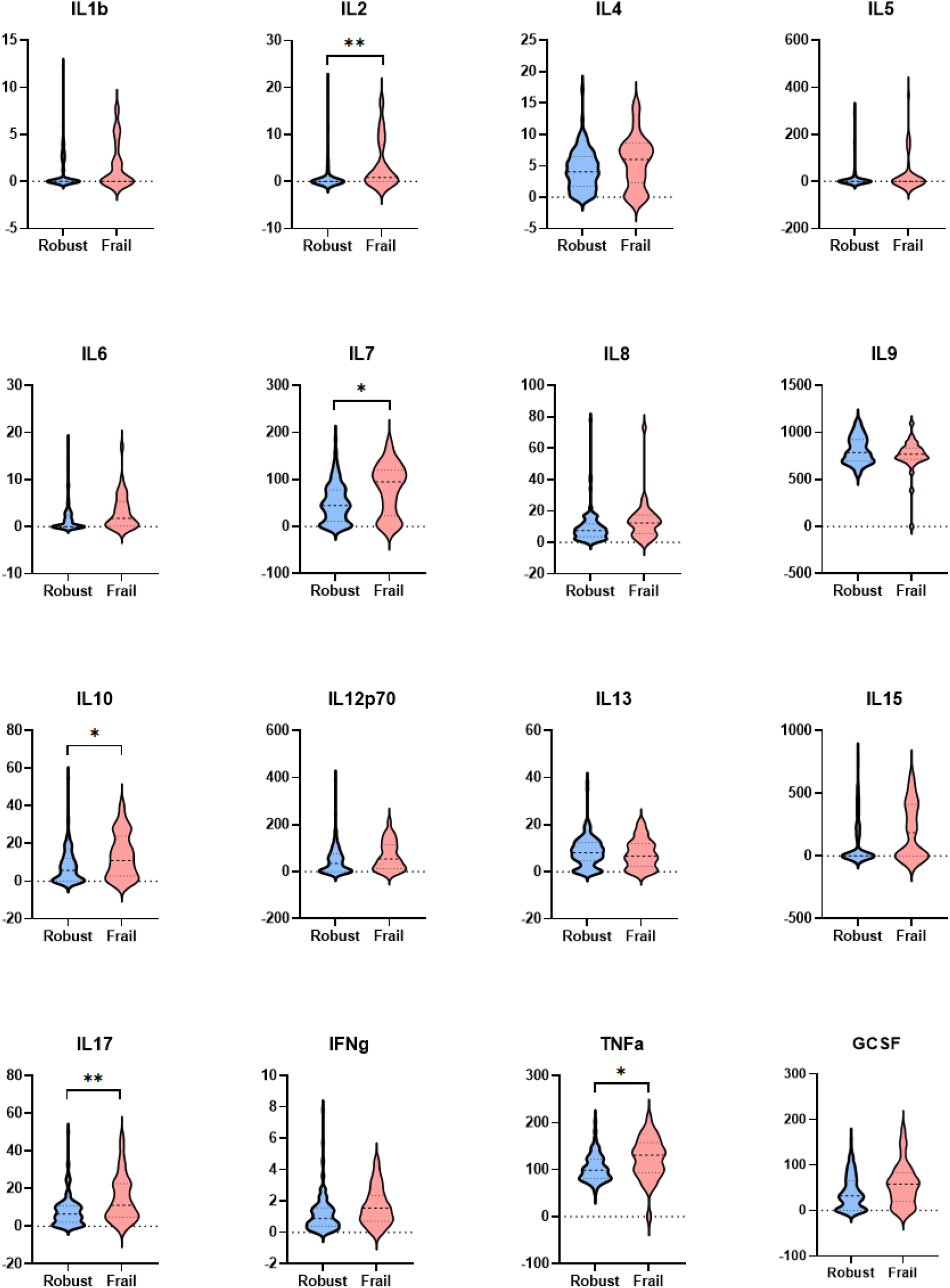
T-regulatory cytokines are altered in frail older adults. Unadjusted violin plots comparing levels of serum inflammatory markers between robust and frail participants. *p<0.05, **p<0.01 using Holm-Šídák method for multiple comparisons.

To account for differences in baseline characteristics between the two populations, we used logistic regression analysis to perform an adjusted analysis, controlling for clinical covariates using stepwise backward elimination. Consistent with our results above, our regression revealed that age, ADAS-Cog 13 scores, and MNA Scores were independent predictors of frailty. Our analysis also revealed that frailty was associated with the presence of hypertension and use of antipsychotic medications (data not shown). After adjustment, three cytokines remained significantly associated with frailty (Table 2 and Supplemental Table 1). Differences in IL-2 were found to have the greatest magnitude of effect on the presence of frailty. TNFa, which was previously noted to differ between groups no longer reached the threshold for significance following adjustment (Table S1).

**Table 2.** Odds ratios of frailty for unadjusted and logistic regression adjusted inflammatory marker values. Brackets represent 95% confidence interval. FDR, false discovery rate.

|  | Unadjusted OR | Adjusted OR | p | FDR |
| --- | --- | --- | --- | --- |
| IL2 | 1.15 [1.06 - 1.27] | 1.17 [1.06 - 1.30] | <0.01 | 0.03 |
| IL10 | 1.06 [1.02 - 1.10] | 1.08 [1.03 - 1.14] | <0.01 | 0.03 |
| IL17 | 1.07 [1.03 - 1.12] | 1.07 [1.02 - 1.12] | <0.01 | 0.04 |

### Inflammatory markers are predictive of frailty

We next sought to develop a predictive model of frailty incorporating blood inflammatory markers and clinical data. To do this, we performed five XGBoost classification experiments using five random seeds, treating frailty as a binary variable. As shown in Figure 3A and 3B, our model demonstrated good sensitivity and specificity (66.67% and 74.10%, respectively) with an area under the curve (AUC) of 0.78. To better assess individual components within the model, we utilized SHapley Additive Explanation (SHAP), a game theory model of feature contributions to the overall machine learning model^27^. Cytokines were the most frequently represented biological category among the top predictors (n=16) and included the single most influential feature, IL7, where higher values strongly contributed to positive model predictions. While growth factors (n=4) showed the highest average predictive importance as a category, the SHAP summary plot indicates they often behaved in the opposite manner; for example, higher levels of the top growth factor, GMCSF, were linked to negative SHAP values. (Fig 3C, 3D, and 3E).

**Figure 3.**
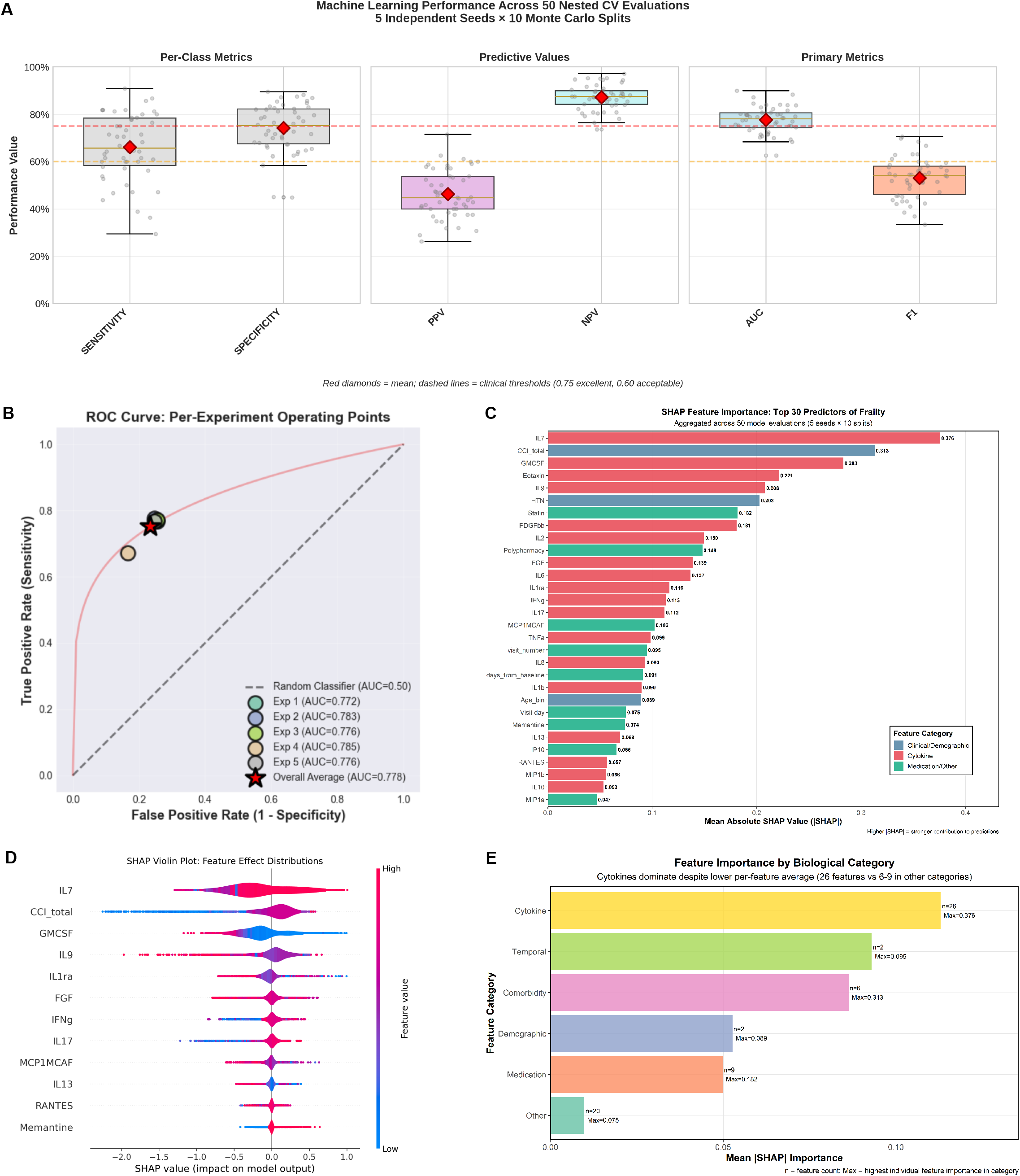
Inflammatory markers are significant predictors of frailty. XGBoost machine learning modeling of frailty incorporating demographics, clinical covariates, and serum inflammatory markers. A) Box and whisker plots of performance characteristics of the model. Dashed red line represents acceptable performance standard. (PPV, positive predictive value; NPV, negative predictive value; AUC, area under the curve). B) Receiver operator curve (ROC) of all model runs. C) SHAP Importance of top 30 predictors of the model. D) Violin plot of SHAP feature effects for individual predictors of the model. E) Feature importance by category based on model performance.

## DISCUSSION

Here we provide evidence of a link between whole-body inflammation and frailty in older adults. Our data indicate that serum inflammatory markers are increased in frail older adults. When utilized in a machine-learning classification model, our data generate an integrated predictive model of frailty which indicated that serum cytokines are the most important feature class in prediction of frailty. Together, these data signify a role of “inflammaging” in the pathogenesis of frailty, and suggest that serum inflammatory may function as an indicator of frailty in older adults.

Our data differ somewhat from prior investigations on inflammation in frailty. A number of prior studies have demonstrated a potential increase in CRP, TNFα and IL-6 in aging and frailty^8,29–32^. While our unadjusted data suggest IL-6 and TNFα were increased in frail patients, following adjustment these did not reach threshold for significance. One potential rationale for this difference is that these markers are associated with parallel syndromes of aging such as cognitive decline or malnutrition which were accounted for in our adjusted analysis. Additionally, several of these studies focused on outcomes which differ from ours, namely all-cause mortality, which, while associated with frailty was not the primary focus of the present analysis^31,32^. In an analysis similar to ours, Mitnitski et al. developed a biomarker model of frailty in which neither IL-6 nor TNFα were significantly influential, though CRP, which was not evaluated in our study, remained an important predictor.

IL-2 had the greatest effect on frailty in our regression analysis, and was demonstrated to have high importance in our machine learning model. Interestingly, IL-2 has been shown to decrease with aging across several mammalian species including humans, which is thought to lead to immunosenescence^33,34^. Consistent with a protective theory of IL-2, low dose IL-2 therapy in mice mitigated effects of a high-fat diet via hypothalamic sympathetic nervous system simulation^35^. Reintroduction of cerebral IL-2 expression also reversed cognitive effects of aging in old mice^36^. Despite this, serum IL-2 was found to be associated with early physiologic aging in obese adults^37^ and increased in adipose tissue insulin resistant individuals^38^, suggesting a role in premature aging and metabolic dysfunction. These discrepancies may indicate temporally specific roles of IL-2 in the process of unhealthy aging. Our cohort was primarily composed of adults with mild frailty (CFS 4 or 5), which reinforces that elevated levels of IL-2 seem to occur early in the disease process and timing of potential interventions may be critical.

Our classification data indicates IL-7 plays an important role in predicting frailty. IL-7 signaling is critical in regulation and maintenance of T-cell populations^39,40^. Consistent with a role in dysfunctional aging, families of “healthy agers” has lower IL-7 receptor expression levels, though nonagenarians themselves had higher overall levels of IL-7 when compared to middle aged controls^41^. Additionally, physical training interventions targeted to reduce frailty in older adults concomintantly reduced levels of IL-7 expression^42^. Contrary to this notion, decreases in IL-7 signaling have been implicated in immunosenescence and diminished lymphocyte in older adults^43^, and recominbant IL-7 has been sought as a potential adjunct in treatment of immune system impairment in severe infection^44,45^. In the context of our data, this could suggest a pleitropic role of IL-7 in aging, and that dysregulation of IL-7 signaling in either direction may lead to maladaptive effects.

Our data implicate regulation of T-cell populations as an important mechanism in the development of frailty. Alterations in T-cell physiology is a known phenomenon of the aging process, coinciding with decreased naive CD8^+^ cells and the presence of unique “age-related” cell types^46^. Studies in aging adults have also demonstrated a decrease in naïve CD4^+^ T-cells, and this has been shown to correlate with frailty and mortality^47,48^. Additionally, markers of T cell senescence are associated with frailty^49^. Taken together with our data, these studies suggest a role for cellular immunity in the pathogenesis of frailty, though the precise signaling mechanisms of this process are not yet clear.

A remaining question is how this pattern of immune regulation might become activated in aging adults. One potential mechanism is via metabolic dysfunction. Adipose tissue from insulin resistant individuals has been shown to have increased levels of IL-2 and immune cell activation^38,50,51^. While not perfectly captured in our data set, lower scores on the Mini Nutritional Assessment in our frail cohort could suggest nutritional intake as a potential source. Another such possibility is intestinal dysbiosis. Gut microbiota undergo age-related changes in composition and has been linked to inflammatory effects^52^. Empiric evidence in older adults reveals dysbiosis is correlated with increased serum inflammatory markers, including IL-2, IL-10, and IL-17^53^. Though the precise mechanisms remain elusive, dysbiosis and metabolic dysfunction may provide upstream targets of inflammation in frailty.

### Strengths and Limitations

Our study has several strengths. First, the GAINS study represents a prospective cohort of community-dwelling older adults with robust demographic and clinical data, allowing for control of potential confounding variables influencing our outcomes. Additionally, we have employed powerful unbiased statistical analysis to generate a model of frailty, which is consistent with a role for immune regulatory molecules in this process. Nonetheless, our study is not without weaknesses. Participants in the GAINS cohort have relatively low frailty scores, limiting extrapolation of our data to older adults suffering from more severe frailty. While we attempted to control for important demographic and clinical variables that may influence frailty, there remains the possibility that additional confounders were not captured in our study, and may influence conclusions. Finally, our data represent a single timepoint in the aging process and development of frailty, which may miss dynamic effects of inflammation and cellular immunity in the aging process.

## Conclusion

In conclusion, we have demonstrated a pattern of inflammatory cytokines in older adults suffering from frailty. In particular, our data implicate T-cell regulation as a predictive and potetnailly causal element in this phenotype. Our classification model demonstrated strong performance in identifying frailty, and that peripheral inflammatory markers were the strongest predictive factors in this determination. Together, these data implicate T-cell regulatory pathways as an important potential mechanism in dysfunctional aging.

## ACKNOWLEDGEMENTS

We would like to acknowledge the staff at the Clinical Research Center at UMass Memorial Medical Center and the Center for Clinical and Translational Sciences at UMass Chan Medical School for clinical facility support during the study period.

## FIGURE LEGENDS

**Supplemental Figure 1.**
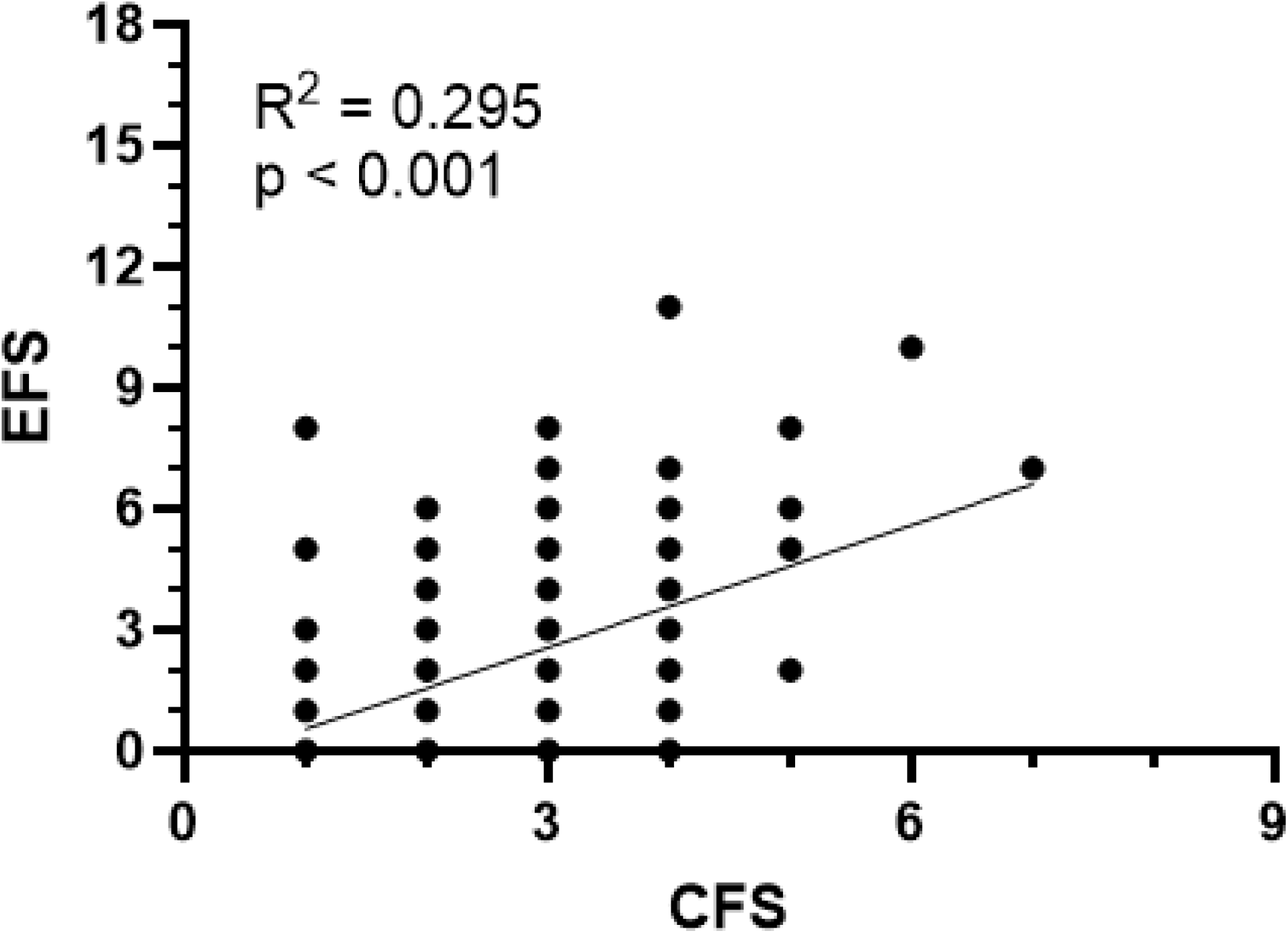
Edmonton Frail Scale versus Clinical Frailty Scale for GAINS Cohort.

**Supplemental Table 1.** Unadjusted and logistic regression adjusted odds ratios (OR) of frailty for serum inflammatory markers.

| Variable | Unadjusted OR | p | Adjusted OR | p | FDR |
| --- | --- | --- | --- | --- | --- |
| IL1b | 1.22 [1.02 - 1.47] | 0.03 | 1.14 [0.90 - 1.42] | 0.25 | 0.37 |
| IL1ra | 1.00 [1.00 - 1.00] | 0.48 | 1.00 [1.00 - 1.00] | 0.22 | 0.36 |
| IL2 | 1.15 [1.06 - 1.27] | <0.01 | 1.17 [1.06 - 1.30] | <0.01 | 0.03 |
| IL4 | 1.12 [1.00 - 1.26] | 0.05 | 1.14 [0.99 - 1.31] | 0.07 | 0.18 |
| IL5 | 1.01 [1.00 - 1.01] | 0.05 | 1.01 [1.00 - 1.02] | 0.09 | 0.19 |
| IL6 | 1.14 [1.03 - 1.28] | 0.01 | 1.10 [0.96 - 1.27] | 0.16 | 0.28 |
| IL7 | 1.01 [1.01 - 1.02] | <0.01 | 1.01 [1.00 - 1.02] | 0.04 | 0.18 |
| IL8 | 1.04 [1.00 - 1.08] | 0.04 | 1.04 [1.00 - 1.08] | 0.07 | 0.18 |
| IL9 | 1.00 [0.99 - 1.00] | 0.04 | 1.00 [0.99 - 1.00] | 0.07 | 0.18 |
| IL10 | 1.06 [1.02 - 1.10] | <0.01 | 1.08 [1.03 - 1.14] | <0.01 | 0.03 |
| IL12p70 | 1.01 [1.00 - 1.01] | 0.08 | 1.01 [1.00 - 1.01] | 0.11 | 0.23 |
| IL13 | 0.98 [0.92 - 1.04] | 0.54 | 1.00 [0.92 - 1.08] | 1.00 | 1.00 |
| IL15 | 1.00 [1.00 - 1.00] | 0.01 | 1.00 [1.00 - 1.01] | 0.03 | 0.15 |
| IL17 | 1.07 [1.03 - 1.12] | <0.01 | 1.07 [1.02 - 1.12] | <0.01 | 0.04 |
| Eotaxin | 1.03 [1.00 - 1.07] | 0.03 | 1.01 [0.97 - 1.05] | 0.71 | 0.81 |
| FGF | 1.02 [0.99 - 1.05] | 0.19 | 1.00 [0.97 - 1.04] | 0.81 | 0.88 |
| GCSF | 1.01 [1.00 - 1.02] | 0.02 | 1.01 [1.00 - 1.03] | 0.02 | 0.12 |
| GMCSF | 1.00 [0.95 - 1.06] | 0.86 | 1.02 [0.96 - 1.09] | 0.52 | 0.66 |
| IFNg | 1.37 [1.01 - 1.86] | 0.04 | 1.35 [0.92 - 2.04] | 0.14 | 0.25 |
| IP10 | 1.00 [1.00 - 1.00] | 0.87 | 1.00 [0.99 - 1.00] | 0.37 | 0.49 |
| MCP1MCAF | 1.05 [0.99 - 1.12] | 0.08 | 1.01 [0.93 - 1.09] | 0.88 | 0.91 |
| MIP1a | 1.29 [0.93 - 1.79] | 0.13 | 1.24 [0.82 - 1.85] | 0.30 | 0.43 |
| RANTES | 1.00 [1.00 - 1.00] | 0.30 | 1.00 [1.00 - 1.00] | 0.60 | 0.72 |
| TNFa | 1.02 [1.01 - 1.03] | <0.01 | 1.01 [1.00 - 1.03] | 0.08 | 0.19 |

